# Spatial distribution and physicochemical properties of respirable volcanic ash from the 16-17 August 2006 Tungurahua eruption (Ecuador), and alveolar epithelium response *in-vitro*

**DOI:** 10.1101/2022.06.21.496955

**Authors:** Julia Eychenne, Lucia Gurioli, David Damby, Corinne Belville, Federica Schiavi, Geoffroy Marceau, Claire Szczepaniak, Christelle Blavignac, Mickael Laumonier, Jean-Luc Le Pennec, Jean-Marie Nedelec, Loïc Blanchon, Vincent Sapin

## Abstract

**Background:** Tungurahua volcano (Ecuador) intermittently emitted ash between 1999 and 2016, enduringly affecting the surrounding rural area and its population, but its health impact remains poorly documented.

**Objectives:** We aim at assessing the respiratory health hazard posed by the 16-17 August 2006 most intense eruptive phase of Tungurahua.

**Methods:** Based on detailed field surveys and grain size analyses, we mapped the spatial distribution of the health-relevant ash size fractions produced by the eruption in the area impacted by ash fallout. We used Scanning Electron Microscopy and Raman Spectroscopy to quantify the mineralogy, composition, surface texture and morphology of a respirable ash sample isolated by aerodynamic separation. The cytotoxicity and pro-inflammatory potential of this respirable ash towards lung tissues was assessed *in-vitro* using A549 alveolar epithelial cells, by Electron Microscopy and biochemical assays (LDH assay, RT-qPCR, multiplex immunoassays).

**Results:** The eruption produced a high amount of inhalable and respirable ash (12.0-0.04 kg/m^2^ of sub-10 µm and 5.3-0.02 kg/m^2^ of sub-4 µm ash deposited). Their abundance and proportion vary greatly across the deposit within the first 20 km from the volcano. The respirable ash is characteristic of an andesitic magma and no crystalline silica is detected. Morphological features and surface textures are complex and highly variable, with few fibres observed. *In-vitro* experiments show that respirable volcanic ash are internalized by A549 cells and processed in the endosomal pathway, causing little cell damage, but some changes in cell morphology and membrane texture. The ash trigger a weak pro-inflammatory response.

**Discussion:** These data provide the first understanding of the respirable ash hazard near Tungurahua, and the extent to which it varies spatially in a fallout deposit. Given the long exposure duration of the surrounding population, the chronic effects of this inhalable, weakly bio-reactive ash on health could be further investigated.

## 1 Introduction

Volcanic eruptions are unpredictable natural phenomena that disperse gaseous species and particles into the atmosphere up to thousands of kilometres away from the source volcano, degrading air quality over vast territories. After the Córdon Caulle eruption in Chile in 2011, for example, increased particulate matter (PM) attributable to volcanic emissions was registered on air quality monitoring stations 1600 km away, in Uruguay.^1^ Closer to active volcanic vents, PM10 concentrations (i.e., PM < 10 µm in size, hence inhalable through the nose and mouth^2^) exceeding the World Health Organization (WHO) 24-hr mean health safety threshold (50 μg/m^3^) are regularly recorded, such as in Iceland.^3,4^ Today, more than 800 million people across the globe live within 100 km of active volcanoes, and 225 million within 30 km, many in developing countries.^5^ Communities benefit from fertilized soil and new rock resources, allowing for the emergence of economic activities such as agriculture, quarrying/mining and tourism. But these populations also frequently breath an ambient air laden with volcanic particles, not only because volcanic activity can be recurrent for long periods of time (e.g., 1995-2013 Soufrière Hills eruption on the Island of Montserrat^6,7^), but also because, after deposition on the ground, volcanic particles are easily remobilized by the wind and human activity.^8,9^

The quality of the air we breathe is a major public health concern in the 21st century. In 2019, more than 6 million premature deaths are attributable to fine-particle pollution worldwide^10^, mainly due to respiratory and cardiovascular diseases. Inhalation of certain mineral dusts, such as asbestos and crystalline silica, is also known to cause acute respiratory diseases and chronic, often irreversible, diseases such as pneumoconioses and cancers.^11^ The comparison to mineral dust is relevant because volcanic emissions are dominated by particles, called “ash”, which are heterogeneous mixtures of crystalline and amorphous silicates (i.e., mineral species with a chemical composition dominated by Si, and rich in Al, Fe, Mg, Na, Ca, K and Ti). Volcanic ash are formed by the fragmentation and quenching of magma erupting at the surface of the Earth, and the erosion of bedrock^12^, and have hence polydisperse size distributions (from 2 mm to < 1 nm), with compositions and physical properties unique to each eruption.

Acute health effects have been documented in exposed populations during several volcanic eruptions (e.g., 1980 Mount St Helens, USA; 1995-2013 Montserrat Soufrière Hills, Caribbean; 1995-today Sakurajima, Japan), including the exacerbation of pre-existing chronic lung diseases.^13,14^ A variable pathogenicity of volcanic ash for the respiratory system has been evidenced by *in-vivo* and *in-vitro* studies. *In-vivo* studies using volcanic ash from the Mount St Helens and Soufrière Hills eruptions indicate inflammation, fibrosis in the lungs^15^, and granuloma in the lymph.^16^ *In-vitro* assays show that volcanic ash can induce an inflammatory response from macrophages and alveolar epithelial cells, despite a limited membranolytic activity.^17-19^ A recent study using sub-4 µm volcanic ash (hence “respirable”; i.e., capable of entering the gas-exchange region of the lungs) from the Soufrière Hills eruption shows that it can activate the well-established NLRP3 inflammatory pathway^20^, potentially due to the high crystalline silica component of this sample (about 15 wt.%^21^). The *in-vitro* response to volcanic ash is generally lower than that observed for crystalline silica^20^, and varies among samples^19^. The physicochemistry of volcanic ash has been studied in relation to the bioreactivity described above, and key properties relevant to toxicity have been identified, namely the particle size distribution, specifically the content in particles finer than 10, 4, 2.5 and 1 µm^22^, the particle morphology^17,23^, the mineralogical (e.g., crystalline silica content) and chemical composition^17,24^, and the oxidative potential of the particles^25^.

Despite this knowledge, anticipating the health hazard at a volcano where no health-focused studies have been carried out remains a challenge. This is due to the high variability of volcanic ash from one volcanic eruption to another, which limits the extent to which the lessons learnt in one volcanic environment can be transferred to another.^14^ The present work aims to characterize health-relevant properties of the volcanic ash from the 1999-2016 Tungurahua eruption, in Ecuador, and to assess the bioreactivity *in-vitro* using alveolar epithelial cells. This long-lasting eruption of Tungurahua has strongly and enduringly affected the surrounding rural area and its population^26^, due to the volcanic emissions, and in particular the large amount of volcanic ash, recurrently dispersed in the environment by explosive episodes.^27-29^ The resuspension of the ash-rich, powdery soils created during the nearly two decades of eruption means that some of the hazards and impacts from this eruption persist even today. Tungurahua is ranked as the most hazardous volcano in South America^30^ due to the high recurrence rate of high-intensity explosive eruptions but little is documented about the impact of the eruptions on population health. Studying the health hazard presented by the activity of this volcano is thus of the utmost relevance. After describing the spatial distribution of inhalable and respirable volcanic ash dispersed in the environment during the most intense explosive phase of the eruption, which occurred on 16-17 August 2006, we describe the physicochemical properties of the respirable ash, and assay the alveolar epithelium response to this ash *in-vitro*.

### 1.1 Background on the Tungurahua social-ecological system

The 5023 m-high Tungurahua volcano is located in the southern part of the Eastern Andean Cordillera, at the limit with the Amazon basin (Fig. 1a). About 20,000 people live in the small tourist town of Baños at the northern foot of the volcano. The lower northwest, west and southwest slopes of the volcano, as well as the western plateau of Quero, are farmed and populated by small rural communities (Fig. 1b, Fig. 2a). A total of 32,000 people live within 15 km of the volcano. Tungurahua is a stratovolcano that began a new explosive eruption at the end of 1999 after 80 years of repose.^28^ This eruption lasted until 2016 and alternated phases of repose, and low to high intensity explosive activity, at the rate of 2 to 3 phases of activity per year on average (Fig. 1c).^31,32^ The magma emitted was andesitic in composition (characterised by about 58 wt.% of SiO^2^). During phases of activity, particle and gas rich volcanic plumes rose 1 to 18 km above the crater and, due to the prevailing East-to-West winds, dispersed volcanic ash on the western and, more rarely, the southern and northern areas.^27-29,33^ The volcano slopes were also regularly affected by pyroclastic density currents (PDCs), which are mixtures of hot fragmented volcanic material and gas that flow down the topography^34^, depositing ash. The most explosive phase of activity (paroxysmal phase) occurred on 16-17 August 2006, lasted about 4 hours, and produced 100.8 ± 21.1 × 10^9^ kg of fragmented material.^35^ About 25 wt.% of this material^35^ dispersed in the western direction from a volcanic plume that rose 16-18 km above the crater, and deposited particles up to 60 km away from the volcano in the northern direction and to distances greater than 100 km towards the west.^27^ About 52 wt.% of the fragmented material was transported in PDCs that covered the western slope of the volcano.^35^

**Fig. 1:**
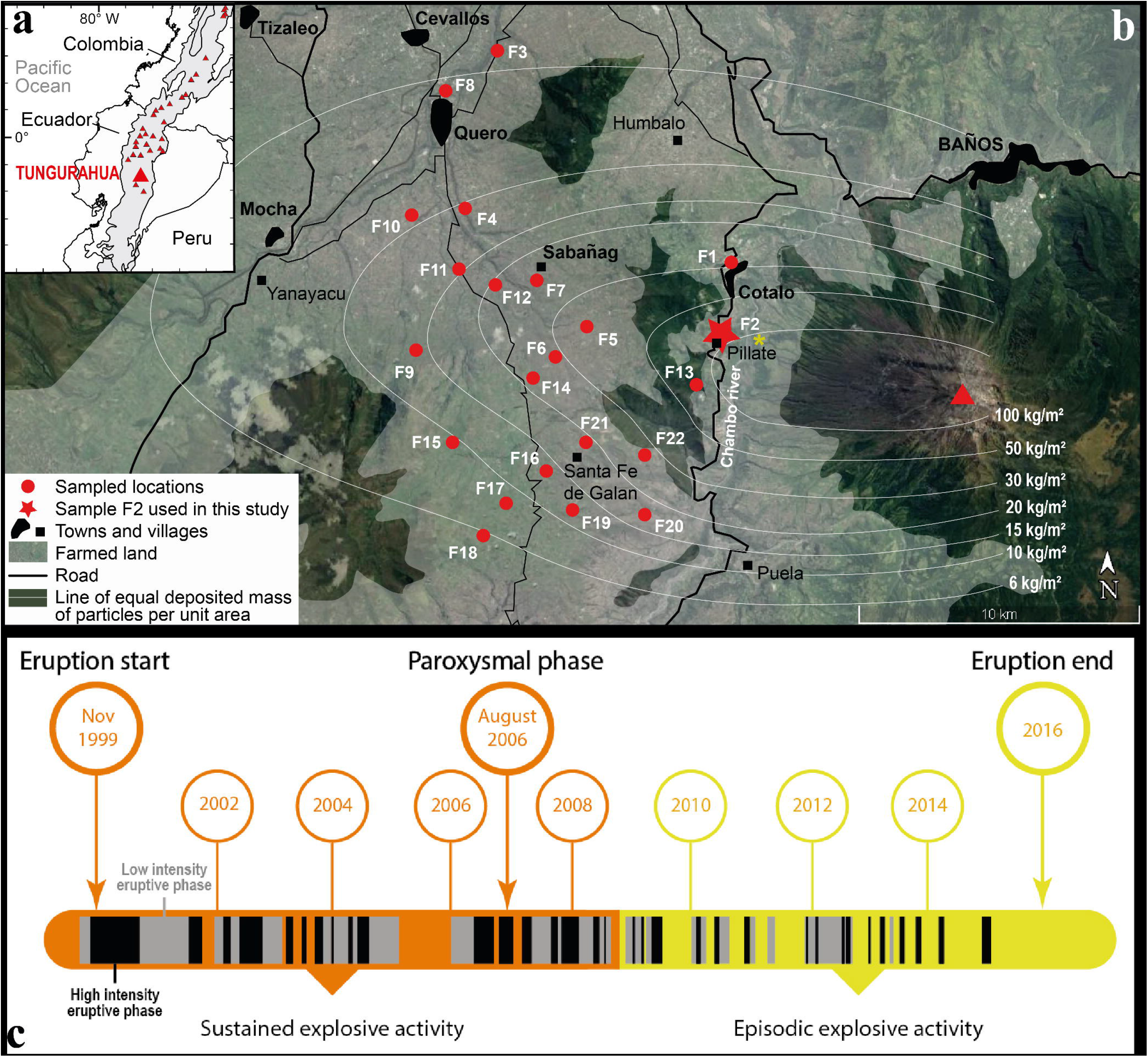
**(a)** Situation map of Ecuador and Tungurahua volcano in the Andean Cordillera (grey shaded area). **(b)** Agricultural area around Tungurahua volcano (red triangle), and locations where the fallout deposit of the 16-17 August 2006 eruptive phase was sampled. The lines of equal deposited mass of particles per unit area during this phase are mapped (white lines), and represent the spatial variations of the amount of particles on the ground. The land in the area is farmed (white transparent area) to the middle of the slope of the Tungurahua volcanic edifice (see also the photograph in Fig. 2a). Location of photograph in Fig. 2b is reported by a yellow star. **(c)** Chronology of the 1999-2016 Tungurahua eruption showing the duration of each explosive eruptive phase and the main changes in intensity (black vs. grey eruptive phases) and steadiness (orange vs. yellow time periods) of the overall activity. Note that the term “eruption” refers to the whole 1999-2016 eruptive period, while “phase” corresponds to the shorter, alternating periods of variably intense explosive activity within the whole duration of the 1999-2016 eruption. Each high (black slots) and low (grey slots) intensity eruptive phase was characterized by explosive plumes reaching an altitude of at least 1 km above the vent and depositing particles in the area defined in **(b)**. After Hidalgo, et al. ^31^ and Muller, et al. ^32^.

**Fig. 2:**
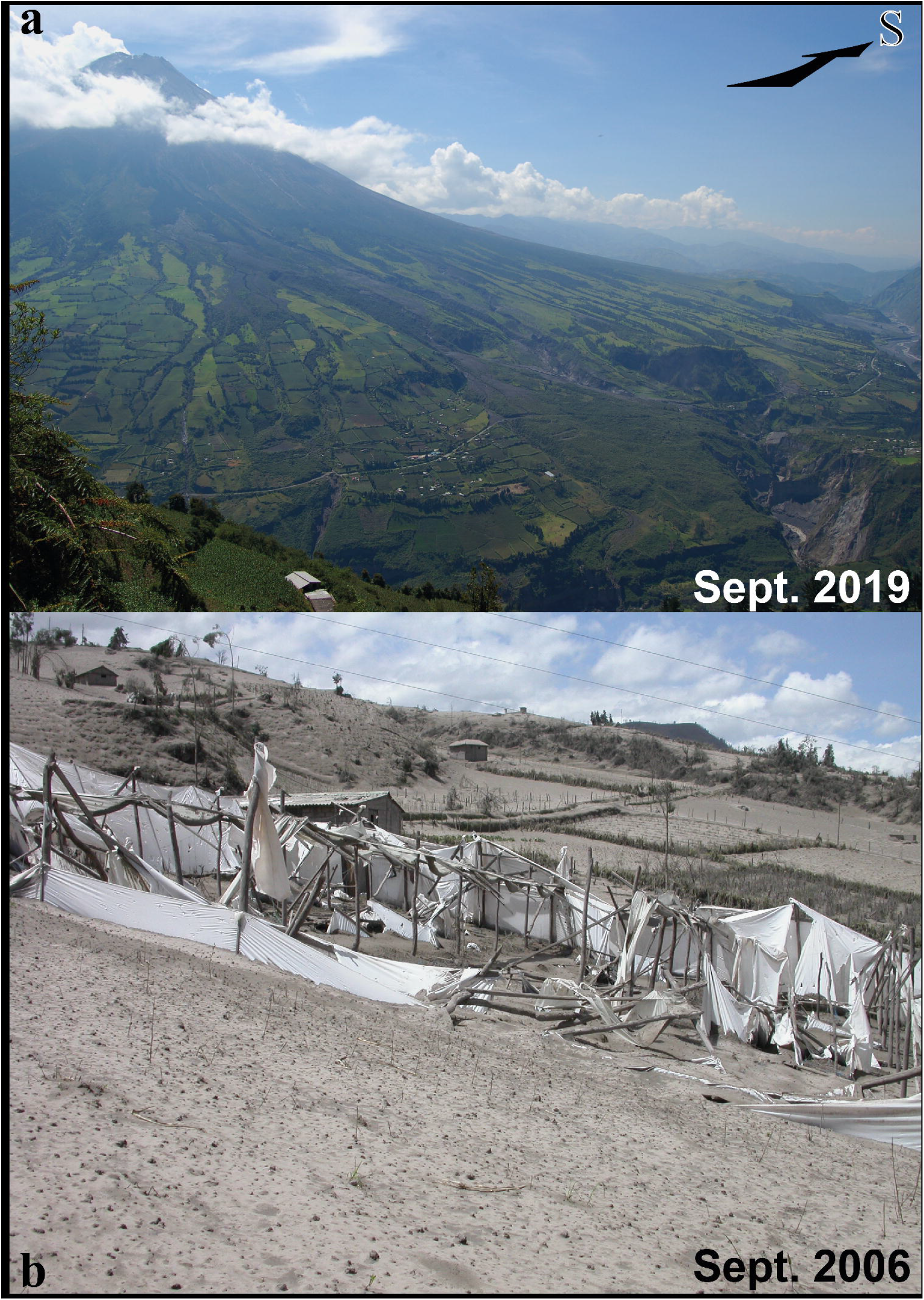
**(a)** Western slope of Tungurahua volcano in September 2019 showing how high the farmed land spreads on the volcano’s flank. Photograph by J. Eychenne taken from location F1 on Fig. 1a. **(b)** Greenhouses and crops covered and destroyed by ash from the explosive plume from the 16-17 August 2006 eruptive phase. Photograph by J.-L. Le Pennec, taken one month after this event. See location on Fig. 1b.

The recurrent eruptive activity between 1999 and 2016 produced powdery deposits (Fig. 2b), which still cover the topography in places around Tungurahua and are continually remobilized by the wind and human activity. They constitute the soils on which human activity has carried on since 1999. Indeed, the local populations were evacuated to nearby areas not affected by the volcanic activity in 1999, but settled back into their homes as soon as 2000.^36,37^ Over the next 20 years, the local people learned how to live with the volcanic risk. They developed a unique crisis management network, based on the close interaction between the scientists in charge of monitoring the volcano and each community.^38,39^ Thanks to voluntary and timely evacuations prior to each of the high intensity explosive phases, the communities were able to remain on their lands throughout the duration of the eruption with few fatalities.^40,41^ The populations coped with the recurring environmental changes caused by the eruptive activity by implementing adaptation strategies that allowed them to minimize the impacts on agriculture.^26,40^ They selected crops resistant to volcanic ash and developed cultivation methods that minimized their vulnerability.^26^ After each major eruptive phase, the crops, meadows, and farms were rapidly cleaned and reclaimed for agriculture. For all of these reasons, Tungurahua is an example of a resilient volcanic social-ecological system. But this remarkable resilience has promoted a prolonged exposure of the population to volcanic PM, exacerbated by the outdoor occupational activity of most people, farming being the principal occupation of 60 to 70% of adults living in the rural surroundings of Tungurahua.^26^ During the 1995-2013 Soufrière Hills eruption on the Island of Montserrat, comprehensive surveys of personal exposure demonstrated that outdoor workers were the most exposed to airborne volcanic ash.^42^ This situation is the motivation to determine the respiratory hazard of the volcanic ash from Tungurahua.

### 1.2 State of knowledge on the health impact of the Tungurahua eruption

Scarce data on the health impact of the long-lasting Tungurahua eruption are available. Analysis of health records up to 2001 from local health care centres and qualitative surveys in the communities surrounding the volcano in 2001 demonstrate an increase in acute respiratory infections and self-reported respiratory problems in the year following the start of the eruption.^43^ Digestive problems related to eating and drinking food and water contaminated by volcanic ash, as well as skin, abdominal and respiratory problems, were reported in 2010, after 10 years of volcanic activity, by healthcare professionals working in local health centres.^44^ Despite these observations, no public health mitigation strategies were implemented to monitor and minimize the impacts of the volcanic eruption on the population. In addition, despite several volcanological studies describing the physicochemistry of the volcanic particles produced by this eruption ^27,45-48^, the respirable ash fraction has never been characterized, neither has their bioreactivity been studied. This work aims at filling these stark gaps in knowledge.

## 2 Material and methods

### 2.1 Mapping the inhalable volcanic ash fractions deposited

During the weeks following the 16-17 August 2006 eruptive phase, the area affected by particle fallout from the explosive plume was surveyed by field campaigns. The fallout deposit generated on the ground was sampled at different locations (Fig. 1b), and the total mass of particles deposited per unit area (MpUA) was measured. The full grain size distribution of each sample was determined by sieving the particles coarser than 63 µm in the laboratory at half phi intervals (where phi=-log^2^ (grain diameter in mm)), and by laser diffraction measurements of the particles finer than 90 µm using a Malvern Mastersizer 2000 (absorption coefficient of 0.1, refractive index of 1.53). Details on field sampling and grainsize analysis methods on this deposit are presented in Eychenne, et al. ^27^. The weight proportions of particles in the different fractions of the inhalable size range (<10, <4, <2.5, <1 µm) were quantified, and converted to mass per unit area of particles deposited using the total MpUA data. These MpUA data for each inhalable volcanic ash fraction were used (1) to quantify their decay rate away from the volcano, and (2) to build maps of their distribution on the ground in the surrounding environment of the volcano, by interpolating manually in-between sampled locations.

### 2.2 Volcanic sample preparation and characterization

#### 2.2.1 Isolation of a respirable ash sample

We selected one key fallout sample from the 16-17 August 2006 eruptive phase, collected on 14 September 2006 in Pillate (Fig. 1b), located 7.9 km away from the eruptive vent along the main axis of plume dispersion and PDC propagation. A respirable fraction (sizes < 4 µm) was physically isolated from this bulk sample using an aerodynamic separation method set up at the USGS California Volcano Observatory, given that sieving to such small grain size is not feasible. In short, the bulk sample was sucked at a steady flow rate equivalent to the terminal settling velocity of 4 µm particles, and allowed to fall in a settling chamber to remove aggregates and particles coarser than about 4 µm. The respirable ash was then collected on a filter holder downstream of the settling chamber.^49^ The grain size distribution of the isolated respirable ash was measured by laser diffraction using a Malvern Mastersizer 3000 at *Institut de Chimie de Clermont-Ferrand*, France (absorption coefficient of 0.1, refractive index of 1.63).

#### 2.2.2 Physicochemical characterization of the respirable ash

The mineralogy of the respirable ash was assessed by Raman spectroscopy at *Laboratoire Magmas et Volcans* (LMV, Clermont-Ferrand, France) on unpolished mounts of grains dispersed on carbon sticky tape, and on polished mounts of grains laid in a light-cured universal micro hybrid composite (dental paste Kentfil+). The spectra were collected on individual grains using an InVia confocal Raman micro-spectrometer (*Renishaw*) and equipped with a 532⍰nm diode laser (200⍰mW output power), a Peltier-cooled CCD detector of 1040 × 256 pixels, a motorised XYZ stage and a Leica DM 2500⍰M optical microscope. The scattered light was collected using a back-scattered geometry. The analytical settings used were a laser power at the grain surface between 0.1 and 1 mW, an acquisition time between 15 and 60 s, a grating of 2400 grooves mm^−1^, a 100x microscope objective and a 20-μm slit aperture (high confocality setting). The wavelength was systematically calibrated prior to analysis, based on the 520.5⍰cm^−1^ peak of Si. Spectra were recorded in the wavenumber range 60-1410⍰cm^−1^ (vibrational frequencies of mineral phases and alumino-silicate network domain of glasses), and occasionally in the 2800-3900⍰cm^−1^ range to check for H^2^O and OH molecules. Individual spectra were interpreted in terms of phase (mineral, glass or mixture of mineral(s) and glass) by fitting the main peaks or bands, using reference libraries (RRUFF™ project and Thermofisher Grams Spectral ID®) and publications^50,51^. The proportion of each of the identified phases in the sample was quantified by acquiring a total of 133 spectra on individual grains, allowing the identification of the mineralogical assemblage of the respirable ash sample.

The morphology, surface texture and composition of the respirable ash were assessed by the *Helios 5* (*ThermoFisher Scientific*) scanning electron microscope coupled with a focused ion beam (Xe plasma; FIB-SEM) at LMV, France, on unpolished mounts of grains dispersed on polycarbonate membranes, stuck on carbon sticky tape and carbon coated. High-resolution images were acquired in secondary electron (SE) and backscattered electron (BSE) modes using electron acceleration voltages of 2 to 5 kV, a current of 50 pA and a 5.0 mm working distance. Elemental composition was measured by energy dispersive X-ray spectroscopy (EDS) with a 60 mm^2^ annular *FLATQUAD* detector (*Bruker*) with beam conditions of 10 kV / 0.8 nA and a 13 mm working distance.

### 2.3 In-vitro bioreactivity assays

#### 2.3.1 Cell culture and treatment with respirable volcanic ash

Immortalized human type II alveolar epithelial cells from the A549 cell line were treated with the isolated respirable volcanic ash sample from Tungurahua. The cells were maintained in culture at 37 °C in a 5% CO_2_ environment in DMEM growth media (Gibco, Thermo Fisher Scientific, Grand Island, USA) supplemented with 10% FBS (Eurobio Scientific, Les Ulis, France), 4 mM L-glutamine (Gibco), and 100 U/ml penicillin, 0.1 mg/ml streptomycin, 0.25 µg/ml amphotericin B (Eurobio Scientific). Twenty-four hours before treatment, cells were trypsinized using a 0.25% solution (Gibco), seeded at a density of 1 × 10^6^ cells/well in 1 ml of complete medium in 6-well plates and allowed to adhere overnight. Cells were treated in quadruplet for either 6 or 24 hours, with suspensions of the Tungurahua respirable volcanic ash or a positive particle control (Min-U-Sil quartz < 10 µm, which is a benchmark sample well-known for its cytotoxicity and inflammatory activity^52^). Particles were suspended at 250 and 1000 µg/ml (based on published dose-response curves for volcanic particles^19^) in serum-free DMEM media and vortexed prior to cell treatment with 1 ml of suspension. These particle concentrations are equivalent to 26 and 105 µg/cm^2^ in 6-well plates. Experiments were repeated 6 times.

After 6 or 24 hours of treatment:(i) culture medium was removed from each well and centrifuged at 13,000 rpm for 10 minutes in 1.5 ml Eppendorf tubes to remove cell debris or remaining particles, and 850 µl of supernatant was then collected and stored at -80 °C until further analyses; (ii) cells were washed 3 times in 1 ml of PBS 1x (Eurobio Scientific), then scratched from the bottom of the wells and centrifuged at 13,000 rpm for 10 minutes in 1.5 ml Eppendorf tubes and stored at -20 °C after removing the supernatant.

Equivalent experiments in terms of exposure durations and doses were performed in duplicate by seeding 5 × 10^5^ cells/well in 12 well plates on Thermanox™ coverslips (Thermo Fisher Scientific, Grand Island, USA) for subsequent preparation and imaging by Transmission Electron Microscopy (TEM) and Field Emission Gun (FEG)-SEM.

#### 2.3.2 Cytotoxicity assay

Cytolysis was assessed by lactate dehydrogenase (LDH) assay in merged culture media of 2 replicates of the 6 independent experiments (leaving duplicates for each experiment). Activity of LDH was quantified with an automated enzymatic assay (Vista, Siemens Health Diagnosis, Paris, France), following the manufacturer’s recommendations.

#### 2.3.3 Assessment of the pro-inflammatory response

The pro-inflammatory response was assessed by quantifying the cytokines interleukin (IL)-6, IL-8, IL-1β and tumor necrosis factor (TNF)-α, both at gene transcript and protein levels. Transcript quantification was measured by real-time quantitative Reverse Transcription Polymerase Chain Reaction (RT-qPCR) using the merged cell pellets from two replicates (leaving duplicates for each experiment) in order to recover enough ribonucleic acid (RNA) for each studied condition. Total RNA extraction was performed using the NucleoSpin RNA Mini kit (Macherey-Nagel GmbH, Düren, Germany) according to the manufacturer’s protocol. RNA concentrations in cell extracts were measured with a DS-11FX spectrophotometer (DeNovix Inc, Wilmington, USA). Complementary deoxyribonucleic acid (cDNA) was synthesized by reverse transcription on 1 μg of RNA using the Superscript IV First-Strand Synthesis system (Invitrogen, Thermo Fisher Scientific, Grand Island, USA), following the manufacturer’s instructions. PCR experiments were performed using specific oligonucleotides (Table 1). RT-qPCR was performed using LightCycler® 480 SYBR Green I Master (Roche, Meylan, France). Transcript quantification was performed in duplicate on four independent experiments. Quantification of amplified transcripts was assessed using standard curves, and gene expression was then normalized to the geometric mean of the housekeeping human genes *RPLP0* (36B4) and *RPS17* (acidic ribosomal phosphoprotein P0 and ribosomal protein S17, respectively), as recommended by the MIQE guidelines.^53^

**Table 1.**
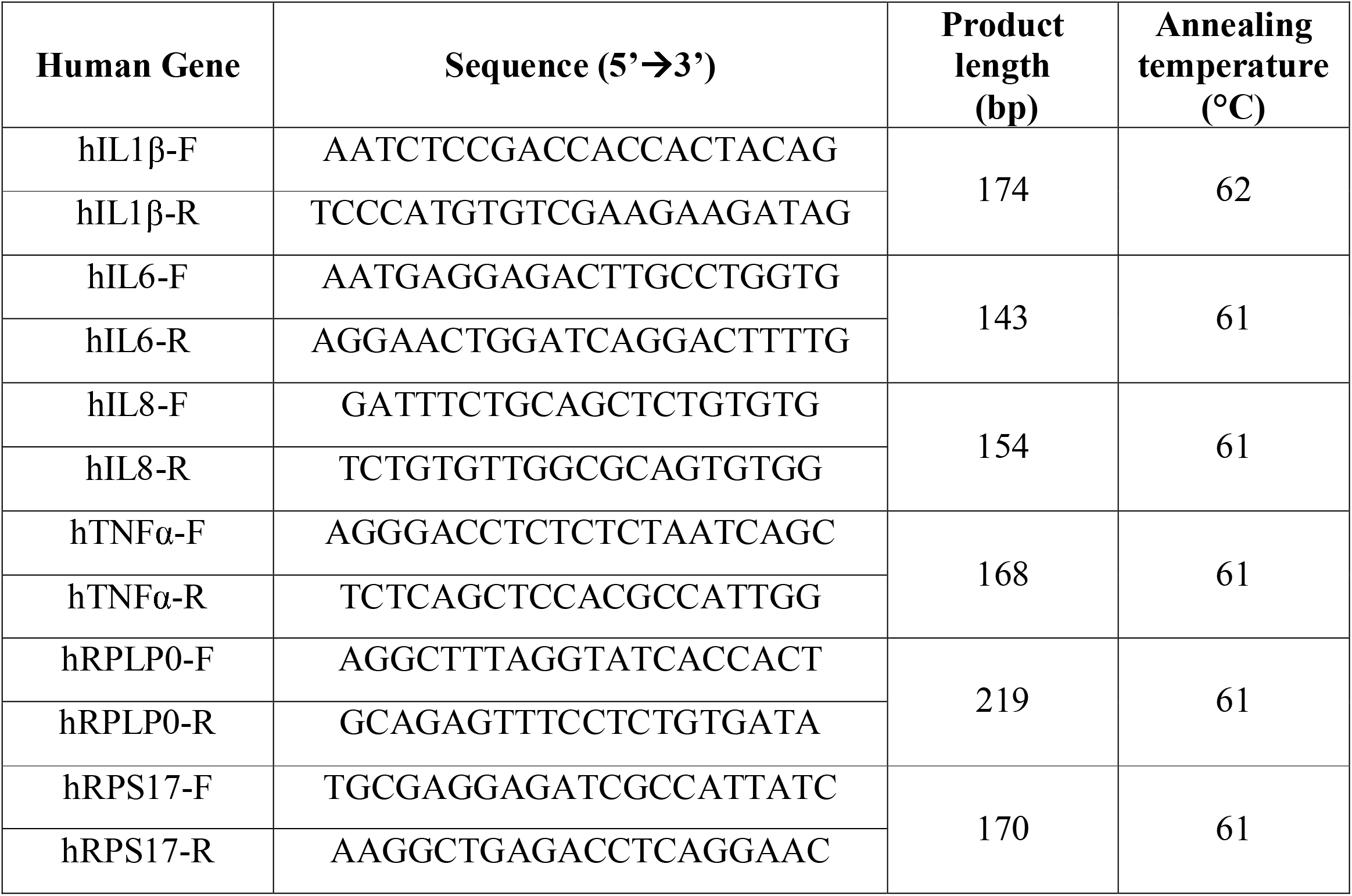
Forward and reverse primer sequences used for RT-PCR and RT-qPCR amplification of human genes.

Cytokine release into the culture media was quantified using automated multiplex immunoassays on Ella™ (San Jose, CA, USA), following the manufacturer’s instructions, on the 4 merged replicates of the 6 independent experiments. Total protein concentrations in the samples were measured using the BCA Protein Assay Kit (Pierce, Thermo Fisher Scientific, Grand Island, USA), and results used to normalize the cytokine concentrations.

#### 2.3.4 Statistical analysis

Statistical analysis was performed using GraphPad PRISM software (San Diego, CA, USA). Results for each condition were compared using the Kruskal-Wallis one-way analysis of variance with Dunn’s post-test. Differences between conditions were considered significant at p ≤ 0.05.

#### 2.3.5 Particle-cell interactions imaging

After 6 or 24 hours of treatment, the cells grown on Thermanox™ coverslips were fixed overnight at 4 °C with 1.6% glutaraldehyde in 0.2 M sodium cacodylate buffer, pH 7.4. For FEG-SEM analyses, samples were then washed for 30 minutes in the same sodium cacodylate buffer (0.2 M, pH 7.4), and post-fixed for 1 hour with 1% osmium tetroxide in the same buffer. After rinsing for 20 minutes in distilled water, dehydration by graded ethanol was performed at 25, 50, 70, 95 and 100% ethanol concentrations (10 minutes at each concentration), to finish in hexamethyldisilazane (HMDS) for 10 minutes. Samples were mounted on stubs using adhesive carbon tabs and coated with chrome (Quorum Q150 TES). Observations were carried out using a Regulus 8230 FEG-SEM (Hitachi, Japan) at 1 and 2 kV with a SE detector and a 13 mm working distance. Elemental microanalysis was performed with an Ultim Max 170 mm^2^ EDS system from Oxford Instrument at 10 kV and a 15 mm working distance.

For TEM analyses, samples were washed three times in sodium cacodylate buffer (0.2 M, pH 7.4), then post-fixed for 1 hour with 1% osmium tetroxide and washed three times (10 min per wash) in this buffer. Samples were dehydrated in a graded ethanol solution (25, 50, 70, 95 and 100% ethanol). They were then infiltrated for 1 hour with each of the following mixtures:(1) 2 volumes of ethanol 100% with 1 volume of EPON resin, (2) 1 volume of ethanol 100% with 1 volume of EPON resin, and (3) 1 volume of ethanol 100% with 2 volumes of EPON resin. Finally, samples were infiltrated with resin overnight at room temperature and polymerized for 2 days in a 60 °C oven. Ultrathin sections (70 nm thick) were cut in the samples using a UC7 ultramicrotome (Leica, Wetzlar, Germany) and stained with uranyl acetate and lead citrate. Note that the hardness contrast between the mineral particles (ash and quartz) and the cells led to tears in some sections. The sample sections were imaged at 80 kV with a TEM H-7650 (Hitachi, Japan) and a Hamamatsu AMT40 camera.

Electron microscopy preparations and analyses were all performed by the *Centre Imagerie Cellulaire Santé* (Clermont-Ferrand, France). All the chemical products were from Electron Microscopy Science, and distributed in France by Delta Microscopies.

## 3 Results

### 3.1 Spatial distribution of the inhalable volcanic ash fractions

Within less than 20 km from the volcano, the fallout deposit from the 16-17 August 2006 paroxysmal phase of the Tungurahua eruption was rich in inhalable particles, with sub-10 µm ash constituting 12.0 to 1.5 wt.% of the deposit, sub-4 µm 6.0 to 0.6 wt.%, sub-2.5 µm 4.8 to 0.5 wt.% and sub-1 µm 2.4 to 0.3 wt.% (Table 2). When normalized to the total mass of particles deposited, these values correspond to 12.0 to 0.04 kg/m^2^ of sub-10 µm ash deposited, 5.3 to 0.02 kg/m^2^ of sub-4 µm, 4.1 to 0.01 kg/m^2^ of sub-2.5 µm and 1.9 to 0.01 kg/m^2^ of sub-1 µm (Table 2).

**Table 2.** Sample list, coordinates (WGS 84 map datum), and sedimentological characteristics (including proportions and mass per unit area (MpUA) of the inhalable volcanic ash fractions). Sample locations in Fig. 3.

| Sample | UTM coordinates<br>(zone 17,<br>hemisphere S) |  | Dist.<br>from<br>vent<br>(km) | Total<br>MpUA<br>(kg/m<sup>2</sup>) | wt.% |  |  |  |  | MpUA (kg/m<sup>2</sup>) |  |  |  |  |
| --- | --- | --- | --- | --- | --- | --- | --- | --- | --- | --- | --- | --- | --- | --- |
|  | E | N |  |  | < 63<br>µm | < 10<br>µm | < 4<br>µm | < 2.5<br>µm | < 1<br>µm | < 63<br>µm | < 10<br>µm | < 4<br>µm | < 2.5<br>µm | < 1<br>µm |
| F 1 | 776681 | 9841976 | 8.5 | 29.1 | 14.4 | 3.7 | 1.5 | 1.1 | 0.4 | 4.2 | 1.1 | 0.4 | 0.3 | 0.1 |
| F 2 | 776416 | 9839600 | 7.9 | 100.2 | 38.8 | 11.9 | 5.3 | 4.1 | 1.9 | 38.9 | 12.0 | 5.3 | 4.1 | 1.9 |
| F 3 | 768261 | 9849616 | 20.6 | 2.7 | 5.9 | 1.5 | 0.6 | 0.5 | 0.3 | 0.2 | 0.04 | 0.02 | 0.01 | 0.01 |
| F 4 | 767139 | 9843956 | 18.1 | 10.3 | 24.2 | 6.2 | 3.0 | 2.4 | 1.3 | 2.5 | 0.6 | 0.3 | 0.2 | 0.1 |
| F 5 | 771469 | 9839734 | 12.8 | 37.2 | 32.3 | 9.6 | 4.6 | 3.7 | 2.1 | 12.0 | 3.6 | 1.7 | 1.4 | 0.8 |
| F 6 | 770323 | 9838700 | 13.9 | 27.0 | 31.7 | 9.3 | 4.5 | 3.6 | 2.0 | 8.6 | 2.5 | 1.2 | 1.0 | 0.5 |
| F 7 | 769701 | 9841314 | 14.9 | 25.9 | 33.9 | 10.5 | 5.1 | 4.1 | 2.1 | 8.8 | 2.7 | 1.3 | 1.1 | 0.5 |
| F 8 | 766408 | 9848198 | 20.6 | 6.3 | 21.2 | 5.2 | 2.2 | 1.8 | 0.8 | 1.3 | 0.3 | 0.1 | 0.1 | 0.05 |
| F 9 | 765393 | 9838940 | 18.8 | 14.4 | 29.0 | 8.3 | 4.1 | 3.4 | 1.8 | 4.2 | 1.2 | 0.6 | 0.5 | 0.3 |
| F 10 | 765203 | 9843728 | 19.9 | 6.7 | 23.1 | 7.6 | 3.7 | 2.9 | 1.5 | 1.5 | 0.5 | 0.2 | 0.2 | 0.1 |
| F 11 | 766895 | 9841782 | 17.7 | 13.8 | 29.9 | 10.1 | 4.9 | 3.9 | 2.0 | 4.1 | 1.4 | 0.7 | 0.5 | 0.3 |
| F 12 | 768190 | 9841254 | 16.3 | 20.0 | 33.7 | 10.3 | 5.0 | 4.1 | 2.2 | 6.8 | 2.1 | 1.0 | 0.8 | 0.4 |
| F 13 | 775412 | 9837718 | 8.8 | 63.5 | 37.7 | 11.8 | 5.9 | 4.8 | 2.4 | 24.0 | 7.5 | 3.8 | 3.0 | 1.5 |
| F 14 | 769533 | 9837938 | 14.7 | 22.4 | 29.7 | 9.1 | 4.3 | 3.4 | 1.8 | 6.7 | 2.0 | 1.0 | 0.8 | 0.4 |
| F 15 | 766750 | 9835702 | 17.6 | 7.8 | 22.1 | 6.5 | 2.8 | 2.2 | 1.2 | 1.7 | 0.5 | 0.2 | 0.2 | 0.1 |
| F 16 | 770009 | 9834660 | 14.6 | 13.4 | 15.5 | 4.1 | 1.9 | 1.5 | 0.8 | 2.1 | 0.5 | 0.3 | 0.2 | 0.1 |
| F 17 | 768594 | 9833600 | 16.2 | 7.0 | 19.1 | 5.6 | 2.5 | 2.0 | 1.0 | 1.3 | 0.4 | 0.2 | 0.1 | 0.1 |
| F 18 | 767830 | 9832468 | 17.3 | 5.0 | 18.8 | 4.3 | 1.9 | 1.5 | 0.8 | 0.9 | 0.2 | 0.1 | 0.1 | 0.04 |
| F 19 | 770958 | 9833310 | 14.1 | 9.3 | 11.2 | 3.1 | 1.4 | 1.2 | 0.6 | 1.0 | 0.3 | 0.1 | 0.1 | 0.1 |
| F 20 | 773566 | 9833020 | 11.7 | 15.4 | 10.9 | 2.7 | 1.3 | 1.0 | 0.5 | 1.7 | 0.4 | 0.2 | 0.2 | 0.1 |
| F 21 | 771414 | 9835712 | 12.9 | 17.0 | 16.7 | 4.6 | 2.2 | 1.8 | 0.9 | 2.8 | 0.8 | 0.4 | 0.3 | 0.2 |
| F 22 | 773510 | 9835168 | 11.0 | 23.9 | 12.3 | 3.1 | 1.5 | 1.2 | 0.6 | 2.9 | 0.7 | 0.4 | 0.3 | 0.2 |

A high spatial variability is observable, with a general decrease with distance from vent of the amount of inhalable ash deposited (Fig. 3). Two decay trends can be noted:(i) a main, steep trend (shaded areas on Fig. 3a,b) corresponding to the variations along the depositional axis oriented west-northwest, where the highest values of MpUA are found for all the inhalable ash fractions (Fig. 3d-g), (ii) a secondary, shallower trend (open symbols on Fig. 3b) corresponding to areas north and south of the depositional axis, where lower values of MpUA are found but spread across a wider spatial area (Fig. 3d-g). Interestingly, the wt.% and MpUA decay trends reveal some differences. The proportions and MpuA of the inhalable ash fractions both decrease with distance from vent (more slowly for the proportions than the MpUA values, Fig. 3a,b). An increase in wt.% with distance is notable between 15 and 20 km from vent within the secondary trends of all the inhalable ash fractions (Fig. 3a), which is not discernible within the MpUA variations (Fig. 3b). The wt.% of the sub-63 µm ash also show a general decrease along the axis and an increase between 15 and 20 km from vent in the off-axis samples (Fig. 3c). The MpUA variations of the inhalable ash fractions mimic the mass variations in the total deposit and in the ash fraction finer than 63 µm (Fig. 3c), with a break-in-slope around 10 km from vent in the decay trend (Fig. 3b,c).

**Fig. 3:**
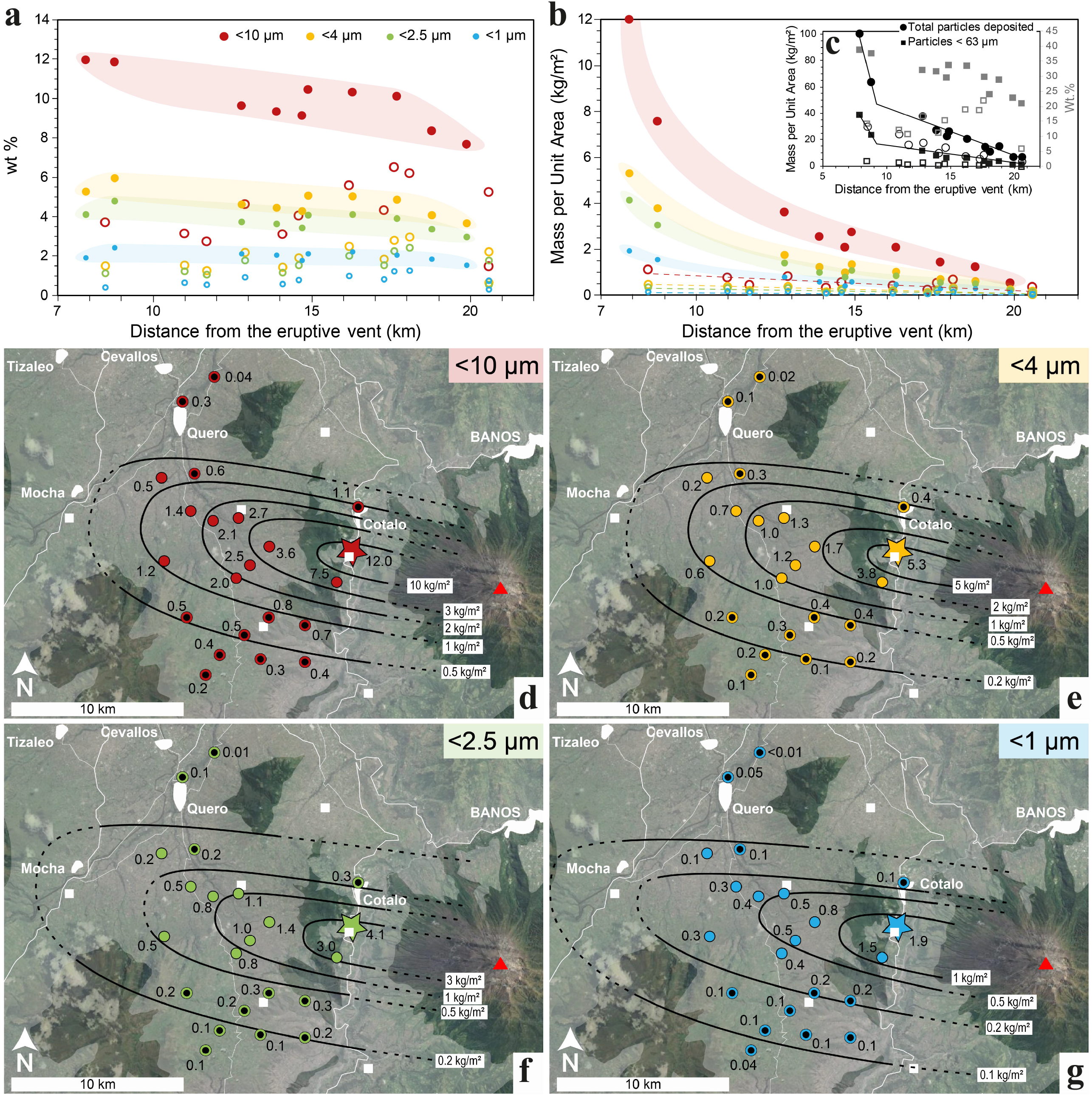
Variations of the inhalable ash fractions across the 16-17 August 2006 Tungurahua fallout deposit. **(a-b)** Decay of deposited sub-10 µm, sub-4 µm, sub-2.5 µm and sub-1 µm volcanic ash with distance from the volcano crater (red triangle on maps; **d-g**) presented as **(a)** mass proportions and **(b)** mass per unit area. Closed and open symbols represent sampled locations on and off the depositional axis, respectively. Colour-shaded areas highlight the on-axis trends. Dashed lines on **(b)** represent the off-axis trends. **(c)** Mass per unit area variations of the bulk deposit and the particles finer than 63 µm with distance from vent in black (left vertical axis). Mass proportions variations of the particles finer than 63 µm with distance from vent in grey (right vertical axis). Closed and open symbols on and off the depositional axis, respectively. The solid lines highlight the decay in mass per unit area along the depositional axis. **(d-g)** Maps of the spatial distribution of the mass per unit area of sub-10 µm **(d)**, sub-4 µm **(e)**, sub-2.5 µm **(f)** and sub-1 µm ash **(g)** in the agricultural area West of Tungurahua volcano, as of September 2006. Closed and open symbols represent on- and off-axis samples, respectively. The solid black lines represent virtual lines of equal deposition (in kg/m^2^) of the considered inhalable fraction (dashed lines extend contours into areas poorly constrained by lack of samples). The white squares are small villages identified in Fig. 1. Coloured star identifies sample used for isolation of respirable ash (F2).

### 3.2 Physicochemical properties of the respirable ash

The grain size distribution (volume %) of the respirable ash sample isolated from sample F2 has a median diameter of 2.5 µm, and a 10^th^ and 90^th^ percentile of 1.1 and 5.9 µm, respectively (Fig. 4a). With 75 vol.% of the sample finer than 4 µm, this ash sample is confirmed respirable.

**Fig. 4:**
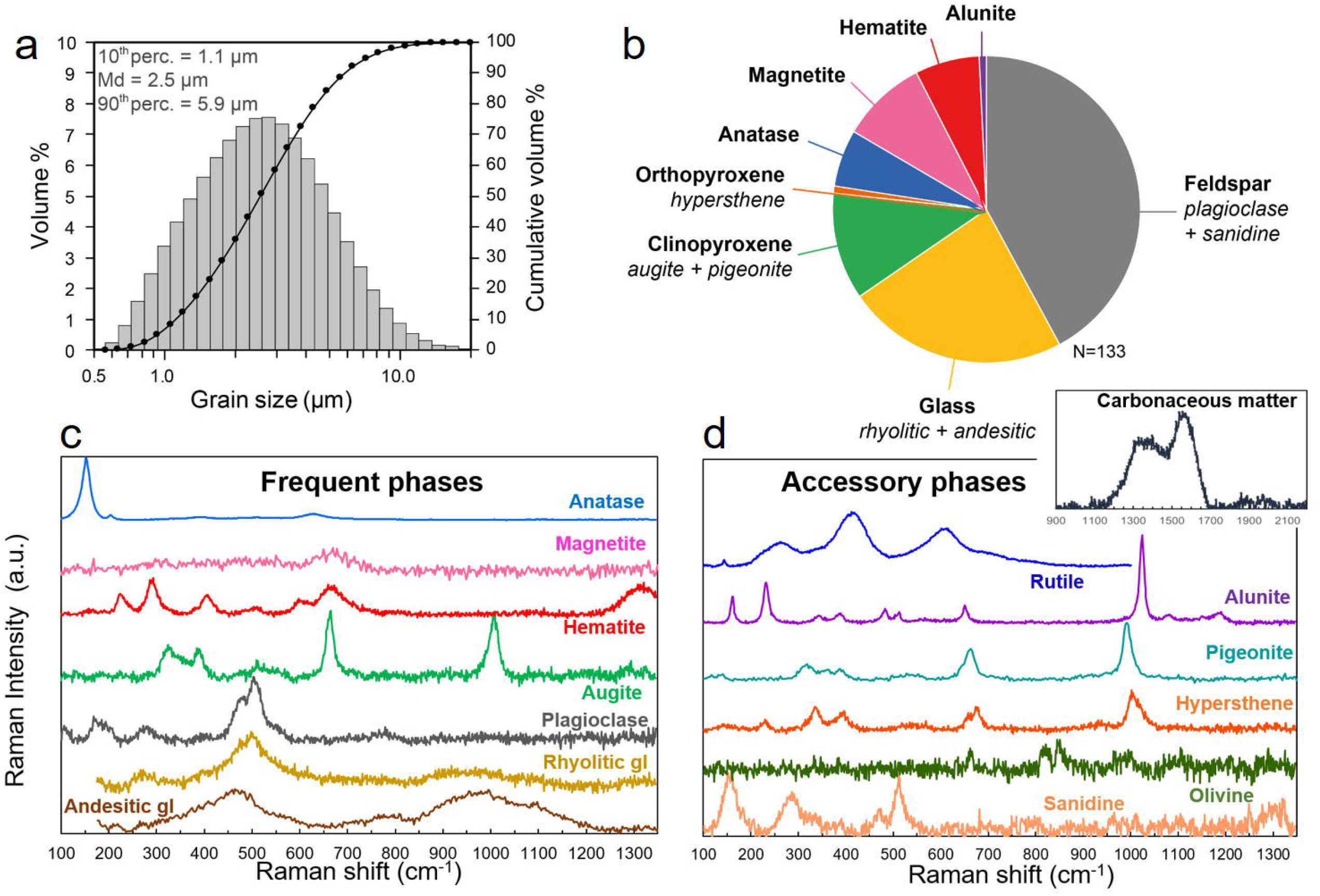
**(a)** Grainsize distribution and **(b-d)** mineralogical componentry of the isolated respirable volcanic ash sample from the August 2006 Tungurahua eruption. **(b)** Number proportion of phases assessed by discrete Raman spectroscopy analyses on 133 individual grains. **(c-d)** Raman spectra for all the mineralogical and carbonaceous phases identified in the sample (vertical axis is arbitrary unit). Accessory phases rutile, olivine and carbonaceous matter each comprise less than 1 nb.% of the sample and are excluded from **(b)**.

The mineralogical phases identified by Raman spectroscopy are:(1) andesitic and rhyolitic glass (constituting 23 nb.% of the phases), (2) crystals of feldspars (plagioclase and sanidine, 42 nb.% of the phases), clinopyroxene (augite and pigeonite, 11 nb.% of the phases), and orthopyroxene (hypersthene, 1 nb.% of the phases), (3) iron and titanium oxides (magnetite is 9 nb.% of the phases, hematite is 7 nb.% of the phases and anatase is 6 nb.% of the phases), and (4) potassium aluminium sulfates (alunite, 1 nb.% of the phases) (Fig. 4b,c and Table 3). Additional accessory phases (less than 1 nb.% of the phases) include rutile, olivine and carbonaceous matter. No crystalline silica was detected by Raman spectroscopy. The standard chemical compositions of the different phases mentioned above are given in Table 3, and confirmed by the SEM-EDS analyses (see chemical maps in Fig. 5).

**Table 3.** Mineralogical phases and number proportions (nb.%) determined by Raman spectroscopy in the respirable volcanic ash sample from the August 2006 Tungurahua eruption. Standard chemical compositions of each identified phase are given for reference. * Glass compositions inferred from microprobe analyses on lapilli from the same eruption in Samaniego, et al. ^45^.

| Phases |  | nb % | Standard chemical compositions |
| --- | --- | --- | --- |
| **Feldspars** | Plagioclase | 42 | Na(AlSi<sub>3</sub>O<sub>8</sub>) - Ca(Al<sub>2</sub>Si<sub>2</sub>O<sub>8</sub>) |
|  | Sanidine |  | K(AlSi<sub>3</sub>O<sub>8</sub>) |
| **Glasses** | Andesitic | 23 | 61.2 ± 0.6 wt.% SiO<sub>2</sub> * |
|  | Rhyolitic |  | 71.6 ± 2.5 wt.% SiO<sub>2</sub> * |
| **Clinopyroxenes** | Augite | 11 | (Ca,Na)(Mg,Fe,Al,Ti)(Si,Al)<sub>2</sub>O<sub>6</sub> |
|  | Pigeonite |  | (Ca,Mg,Fe)(Mg,Fe)Si<sub>2</sub>O<sub>6</sub> |
| **Orthopyroxene** | Hypersthene | 1 | (Mg,Fe)SiO<sub>3</sub> |
| **Anatase** |  | 6 | TiO<sub>2</sub> |
| **Magnetite** |  | 9 | Fe<sup>2+</sup>Fe<sup>3+</sup><sub>2</sub>O<sub>4</sub> |
| **Hematite** |  | 7 | Fe<sub>2</sub>O<sub>3</sub> |
| **Alunite** |  | 1 | KAl<sub>3</sub>(SO<sub>4</sub>)<sub>2</sub>(OH)<sub>6</sub> |
| **Rutile** |  | - | TiO<sub>2</sub> |
| **Olivine** |  | - | (Mg,Fe)<sub>2</sub>SiO<sub>4</sub> |
| **Carbonaceous matter** |  | - | C-C and C-H bonds |

**Fig. 5:**
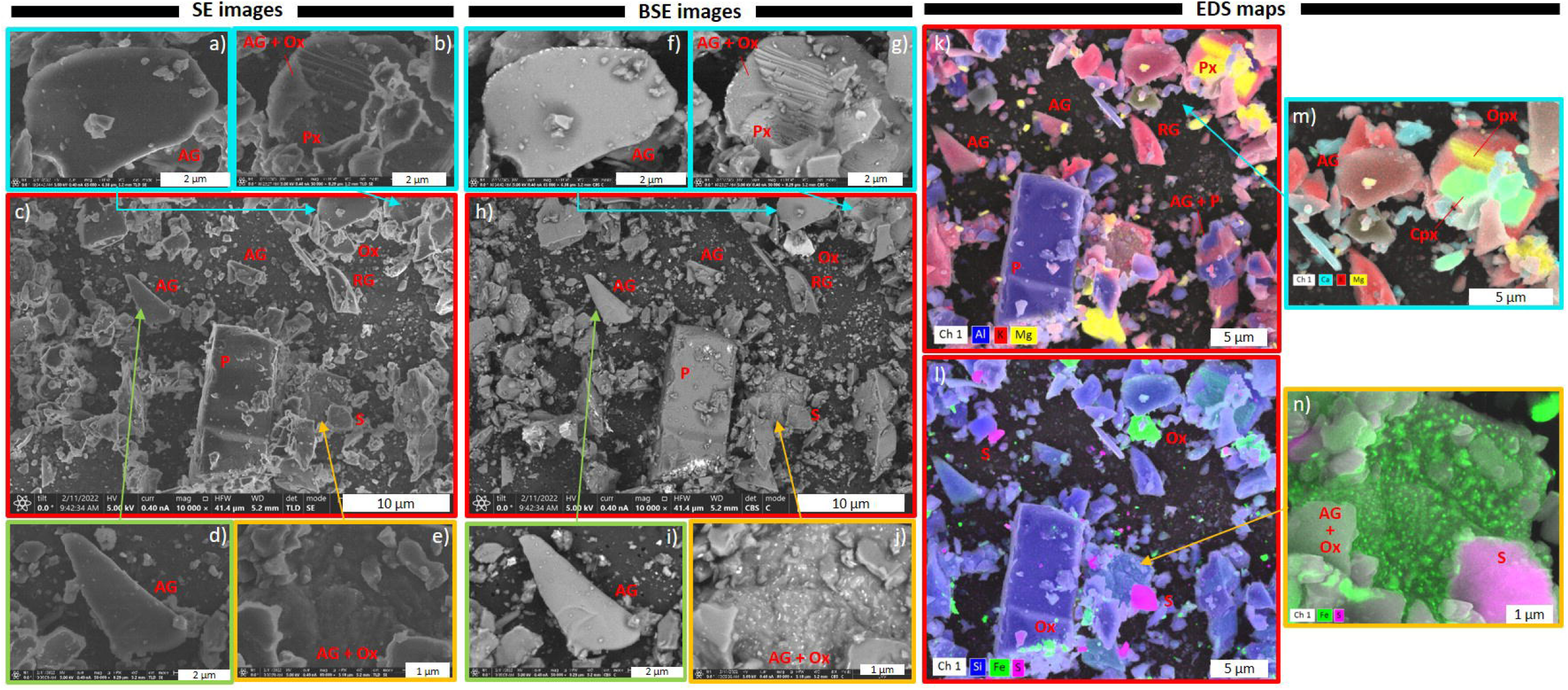
Morphology, texture and surface chemistry of the respirable Tungurahua volcanic ash sample, acquired by FIB-SEM in **(a-e)** SE, **(b-j)** BSE and **(k-n)** EDS modes. AG = andesitic glass; RG = rhyolitic glass; P = plagioclase; Px = pyroxene; Cpx = clinopyroxene; Opx = orthopyroxene; Ox = Fe and Ti oxides; S = Potassium and aluminium sulfates (alunite).

The morphology and surface texture of the particles imaged by SEM are highly variable, and dependent on particle composition (Fig. 5). Glass fragments (K-rich phases on EDS maps, Fig. 5k-n) tend to have concave flat shapes reminiscent of broken vesicle walls, and smooth surfaces (Fig. 5a,c,d,f,h,i). Plagioclase (Al-rich phases on EDS maps, Fig. 5k-n) and pyroxene crystals (Mg-rich on EDS maps, Fig. k-n) are either euhedral, free fragments, or included in glass (Fig. 5b,c,g,h). Clinopyroxene (Ca-rich phases on EDS maps, Fig. 5m) and orthopyroxene are sometime observed associated (Fig. 5m). Crystal sizes vary between > 10 µm in length (euhedral) to < 1 µm (Fig. 5h). Pyroxene crystals tend to be smaller than plagioclase ones. Fractured surfaces are prevalent, both on crystals (Fig. 5b,g) and glass particles. Many Fe-Ti oxides are observed as free < 500 nm particles, or as nanolites in andesitic glass, creating a granular surface (Fig. 5e,j,l,n). The sulfate salts (alunite) are found as free particles ∼3 µm to a few 100s nm in size (Fig. 5l,n). All micrometer-sized particles are covered with smaller particles that are up to a few 100s nm in size. These small particles are either fragments from the andesitic and rhyolitic magmatic assemblage (e.g., glass, pyroxenes, plagioclases, Fe-Ti oxides), or sulfate salts, also visible as small nanometer-sized droplets at higher magnification on glassy surfaces. Very few fiber-like particles are observed.

### 3.3 In-vitro reactivity of the respirable ash

Treatment of A549 cells with the respirable volcanic ash and quartz particles leads to changes in cell morphology, membrane texture and intracellular characteristics (Fig. 6). FEG-SEM images of the A549 cells show that untreated cells create layers of joint cells with spread out shapes (Fig. 6a-d). The surfaces of the membrane display elongated filaments. After 24 hours, slightly retracted cell morphologies can be noted (Fig. 6c-d). Cells treated with respirable volcanic ash have more retracted shapes than untreated cells, and cell junctions are lost (Fig. 6e-g). After 24 hours of treatment, the membrane’s surface appears rough and micro-perforated (Fig. 6g,h). No major differences can be observed between the 250 and 1000 µg/ml doses (Fig. S1 in Supplemental Material). Particles are observed lying on the surface of the cells (Fig. 6f-g) or entangled in their membrane, with filaments embedding the particles (Fig. 6h). Cells treated with quartz particles are entirely retracted with sphere-like morphologies as early as 6 hours after treatment (Fig. 6i-j). Particles are observed as agglomerates on the cells’ surfaces (Fig. 6k) or entangled in the membrane (Fig. 6l).

**Fig. 6:**
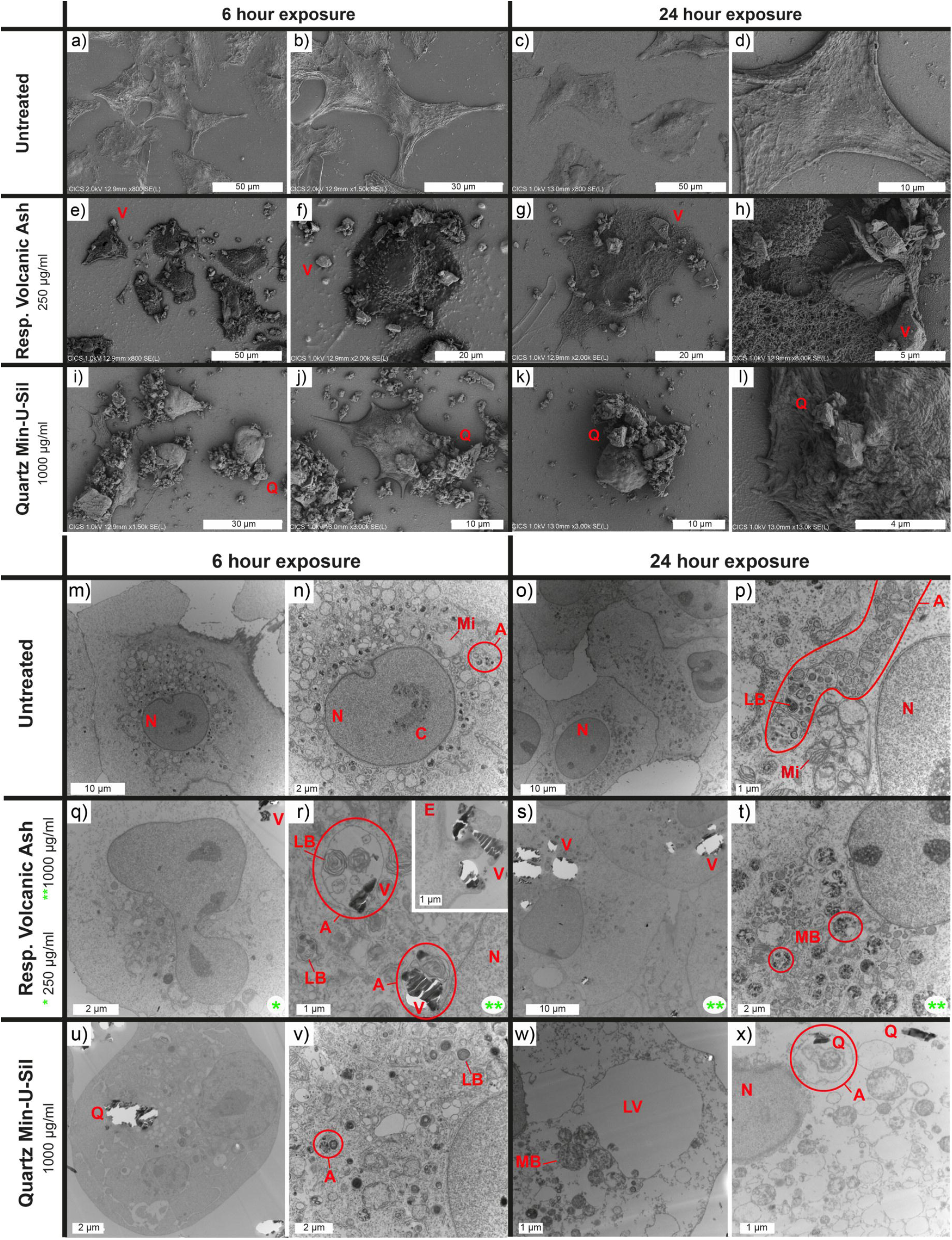
Interactions between A549 cells and respirable Tungurahua volcanic ash or quartz particles imaged by **(a-l)** FEG-SEM and **(m-x)** TEM. **(a-l)** Morphological changes observed after 6 and 24 hours between untreated A549 cells and cells treated with respirable volcanic ash or Min-U-Sil quartz positive control. Images of cells treated with volcanic ash at 250 µg/ml are solely represented here because they offer a better view of the cells and the particles, but no major differences in terms of morphology and surface features are observed between cells treated at 250 and 1000 µg/ml (Fig. S1 in Supplemental Material). Q = quartz particles; V = respirable volcanic ash. **(m-x)** Intracellular changes observed after 6 and 24 hours between untreated A549 cells and cells treated with respirable volcanic ash or quartz Min-U-Sil positive control. Like with the SEM images, no major differences are observed between cells treated with 250 and 1000 µg/ml of volcanic ash (Fig. S2 in Supplemental Material). Green stars indicate treatment dose of volcanic ash (*250 µg/ml, **1000 µg/ml). White bright spots are due to tears in the ultrathin sections, caused by the hardness contrast between the particles and the cells during cutting. A = autophagosomes/autophagolysosomes; C= chromatin; E = endocytosis; LB = lamellar bodies; LV = lysis vacuoles; MB = microvesicular bodies; Mi = mitochondria; N = cell nucleus; Q = quartz particles; V = respirable volcanic ash.

TEM images of A549 cell sections show that untreated cells form a layer with tight junctions and have spread out shapes after 6 and 24 hours of culture in serum-free medium (Fig. 6m,o). Round nuclei containing chromatin can be observed (Fig. 6m-o), as well as numerous organelles in the cytoplasm, including mitochondria and lamellar bodies, the latest producing the pulmonary surfactant in type II epithelial cells (Fig. 6m-p). Many autophagosomes/autophagolysosomes are also present, potentially due to the established upregulated autophagosomal activity of the A549 cell line. Cells treated with respirable volcanic ash contain internalized particles in the cytoplasm, in autophagosomes/autophagolysosomes (where lamellar bodies and mitochondria are often present, too) or multivesicular bodies (Fig. 6q-t). An attempt of endocytosis is also observed at the membrane (Fig. 6r). Despite this upregulated endosomal activity, the cells treated with respirable volcanic ash for 6 and 24 hours preserved their overall integrity (Fig. 6q-t). Cells treated with quartz also contain internalized particles, autophagosomes/autophagolysosomes and microvesicular bodies (Fig. 6u-x). As soon as 6 hours after treatment, cells have retracted shapes and small lysis vacuoles in the cytoplasm (Fig. 6u,v). After 24 hours of treatment, the cells have lost their integrity, with large lysis vacuoles in the cytoplasm, and disintegrated membranes and nuclei (Fig. 6w,x). Both cells treated with respirable volcanic ash and quartz particles contain numerous microvesicular bodies after 24 hours of treatment, which are filled with dark spots (Fig. 6t,w).

LDH activity indicates little effect of the respirable volcanic ash treatment on A549 cell viability (Fig. 7a). Conversely, the quartz treatment triggered a high LDH release, which increased after 24 hours of exposure (Fig. 7a), indicative of time-enhanced cytotoxicity, in agreement with the known toxicity of Min-U-Sil quartz ^52^.

**Fig. 7:**
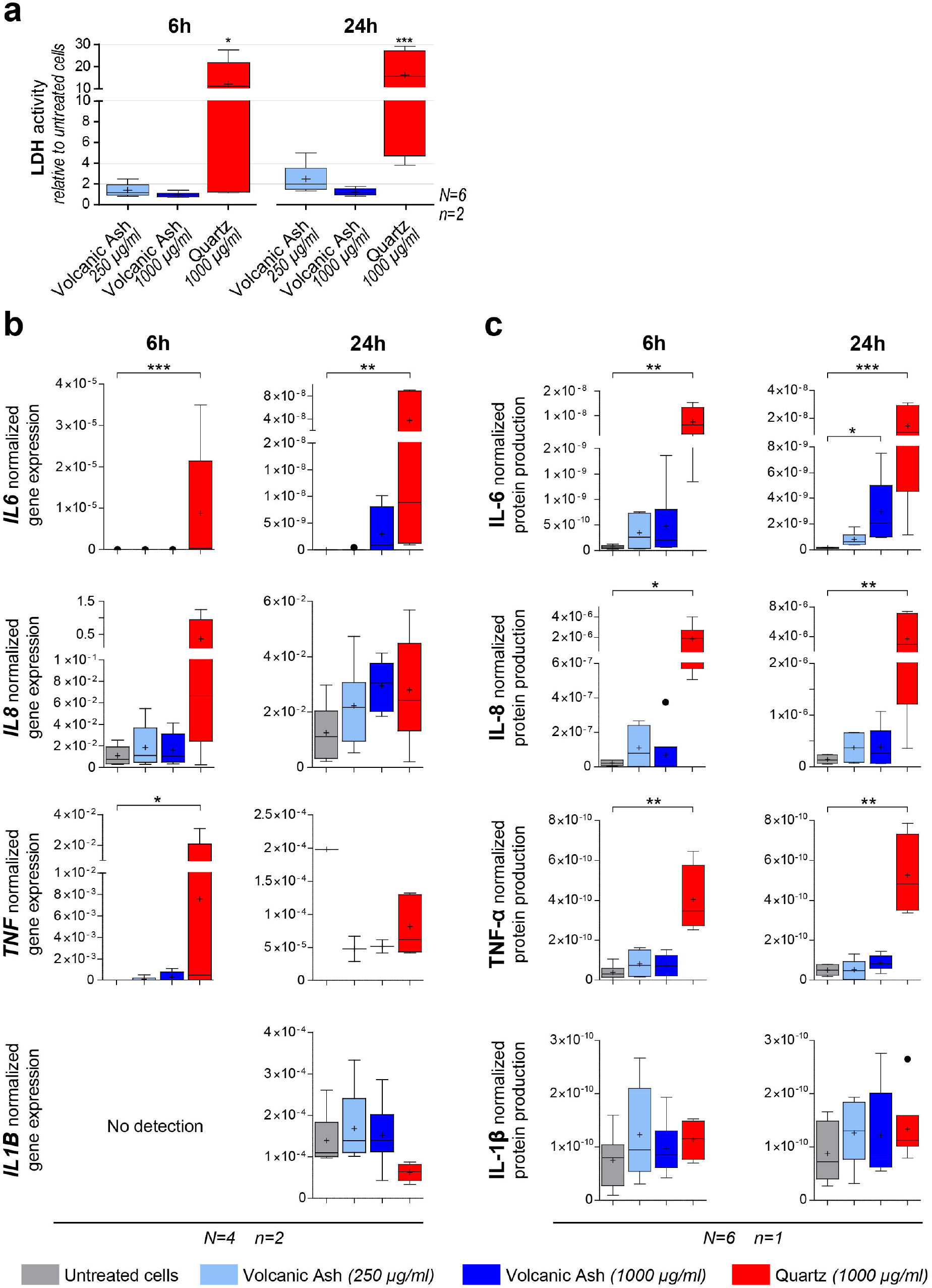
**(a)** Cytotoxicity towards A549 cells of the respirable Tungurahua volcanic ash sample at 2 concentrations and the Min-U-Sil quartz positive control after 6 and 24 hours of exposure, measured by LDH activity relative to untreated cells in duplicates (n=2) of 6 independent experiments (N=6). **(b-c)** Pro-inflammatory response measured by cytokine production by A549 cells after 6 and 24 hours of exposure to 2 concentrations of the respirable Tungurahua volcanic ash and the Min-U-Sil quartz positive control. **(b)** Cytokine gene expression by real-time RT-qPCR on duplicates (n=2) of 4 independent experiments (N=4). Gene expression is normalized to the geometric mean of the expression of two housekeeping genes. **(c)** Cytokine production measured in culture supernatants by automated multiplex immunoassays on Ella™ on the pooled duplicates (for an n=1) of 6 independent experiments (N=6). Protein production presented as the cytokine concentration normalized to the total protein concentration. Boxplots in **(a), (b)** and **(c)** are Tukey boxplots with the mean represented as a cross and the median as an horizontal line. *p ≤ 0.05, **p ≤ 0.01, ***p ≤ 0.001.

The pro-inflammatory response of A549 cells was particle, cytokine, time and dose dependent (Fig. 7b and c). At gene level, a significant upregulation of *IL6* and *TNF* is triggered by the quartz positive control as early as 6 hours after treatment (Fig. 7b). The quartz treatment also augments the expression of *IL8* after 6 hours, but not the expression of *IL1B* (Fig. 7b). After 24 hours of treatment, only the upregulation of *IL6* by quartz is still significant (Fig. 7b). No significant upregulation of *IL6, IL8, TNF* and *IL1B* is detectable after 6 and 24 hours of respirable volcanic ash treatment (Fig. 7b). At protein level, IL-6, IL-8 and TNF-α release was clearly triggered by 6 and 24 hours of quartz treatment (Fig. 7c), while IL-1β production was not triggered by the quartz insult (Fig. 7b and c). A dose- and time-enhanced IL-6 production due to the respirable volcanic ash treatment is observable at the protein level, with a significant difference compared to untreated cells for the high ash dose (1000 µg/ml, or 105 µg/cm^2^) at 24 hours (Fig. 7c). Ash-triggered IL-8 production can also be noted at protein level, but not significantly different from untreated cells. No production of TNF-α or IL-1β occurred after 6 or 24 hours of respirable volcanic ash exposure (Fig. 7c).

## 4 Discussion

### 4.1 Inhalable ash spatial distribution and implications for exposure

The fallout deposit from the paroxysmal explosive phase of 16-17 August 2006 is rich in inhalable and respirable volcanic ash in the studied area (up to 20 km from vent, Fig. 3 and Table 2). A remarkable enrichment in sub-63 µm ash was previously highlighted in this deposit.^27^ It was interpreted as originating from the PDCs emplaced on the volcano’s flanks concomitantly with the rise of the plume at the vent.^27,54^ Sub-63 µm ash escaped by elutriation from the PDCs during flow propagation and generated co-PDC ash clouds that rose in the atmosphere and dispersed towards the West, in the same direction as the main plume formed at the vent. This led to a fallout deposit characterised by a high sub-63 µm ash content^27^, and a decrease in the sub-63 µm ash proportions away from vent along the deposit axis (Fig. 3c). A similar decreasing trend is observed in the proportions of the inhalable ash fractions with distance from vent (Fig. 3a). These trends are counter-intuitive given that all fallout deposits show a decrease in grainsize with distance from vent ^55,56^, which should lead to an increase in the proportions of sub-63 µm and sub-10 µm ash with distance. These unexpected trends are caused by the contribution of elutriated ash discussed above, which decreases with the distance to the PDCs, as demonstrated in Eychenne, et al. ^27^. At Tungurahua, due to the topography, the PDCs always stop in the deep canyon of the Chambo river and remain confined to the East of the Quero plateau (Fig. 1b). These findings indicate that the high amount of inhalable and respirable ash found in the fallout deposit of the August 2006 paroxysmal phase is attributable to the PDC activity. Enrichment of fallout deposits by ash from PDCs is not restricted to the case of the August 2006 Tungurahua eruptive phase, it is a frequent process documented elsewhere (e.g., May 18, 1980 Mount St Helens eruption^57^, 1995-2013 Soufrière Hills eruption^58^). In fact, many eruptions from intermediate to high explosivity produce ash plumes and PDCs concomitantly^35^. Particles transported in PDCs have undergone a secondary fragmentation process (the primary fragmentation being that of the magma in the volcanic conduit) by comminution of bigger grains (grain-to-grain abrasion) during their transport^59-61^. Understanding the origin of the inhalable ash fractions is thus essential, because secondary fragmentation could affect the size, morphology and surface properties of the grains, which are relevant for toxicity.

Importantly, this work highlights a large spatial variability in inhalable ash contents across the fallout deposit (Fig. 3). The variation trends are different depending on whether the inhalable ash content is expressed as proportion or as MpUA. The proportions of the inhalable ash fractions decrease away from vent along the depositional axis, and increase away from vent off-axis. The main MpUA decay trends of the inhalable fractions along the deposit axis are characteristic of fallout deposits, with a break-in-slope around 10 km from vent (notable in the total and sub-63 µm MpUA trends, too; Fig. 3c), reflecting a well-known change in particle settling behaviour from volcanic plumes.^55,62^ Both the proportion and MpUA metrics being based on the deposit-record, they are not representative of what people actually breathe^42^, but of available inhalable and respirable material found in the air during the eruption, and on the ground after particle deposition where they are susceptible to resuspension. Our results establish MpUA of the inhalable ash fractions as a valuable metric for quantifying available material for inhalation, and can support assessment of the respiratory health hazard. Whereas proportions give a relative representation of available ash size fractions relevant for respiratory health, they do not account for the uneven absolute amount of volcanic particles distributed spatially (Fig. 3a). Our work also demonstrates that accounting for the spatial variability in inhalable and respirable ash content is critical for hazard assessment, given that, in our dataset, MpUA values for sub-10 and sub-4 µm ash vary by one order of magnitude within 20 km from the source (Fig. 3b-f). Mapping of the sub-10 µm ash fractions has never been achieved by the volcanology community, which rather focuses on coarser grainsize fractions, more relevant to understanding volcanological phenomena. This makes comparing the inhalable and respirable ash content of the fallout deposit from the 2006 Tungurahua eruption to deposits from other eruptions challenging.

These results have important implications for population exposure around Tungurahua. During the August 2006 eruptive phase, the highest amount of inhalable and respirable ash (up to 12.0 and 5.3 kg/m^2^ of sub-10 and sub-4µm ash, respectively; Table 2) were deposited in the agricultural area of the Quero plateau, where all of the land is farmed and many small villages are located (Fig. 1b). During this eruption, only populations East of the Chambo river (living on the volcano slopes or at the bottom of the edifice; Fig. 1b) were evacuated, and even they eventually settled back on their lands.^40^ The farmers continued to work on their pastures and crops despite the material deposited. The 2006 phase is only one among dozens that dispersed volcanic ash in the western and, more rarely, the southern and northern areas surrounding Tungurahua over the course of the 1999-2016 eruption.^28,29,33^ This means that the amount of inhalable and respirable ash determined in this work represents a minimum estimate. Given that most populations around Tungurahua are outdoor workers^26,40,41^, and that housing and vehicles are permeable due to the mild regional weather, people have a high risk of respiratory exposure to PM in their environment. A similar elevated risk resulting from socio-economic and environmental factors was highlighted during the Soufrière Hills eruption on the island of Montserrat.^6,42^

An important parameter to assess population exposure to volcanic PM is how long ash particles remain available for resuspension in the environment. Some volcanic ash deposits are still remobilised more than a hundred years after eruption, with large regional impacts, e.g., deposits from the 1912 Novarupta eruption in the Katmai region of Alaska.^63^ Timescales of unconsolidated volcanic ash deposit preservation depend on several environmental factors that result in their incorporations and transformation to soils. They include climate, with soils developing slower in drier environments^8,9^, and the nature of the vegetation cover, whereby ash deposits stabilize more efficiently in vegetated environments.^64^ Population exposure to volcanic PM after eruptions is thus likely to be highly variable spatially, and its assessment requires an understanding of the spatial distribution of the ash size fractions relevant for respiratory health within the initial fallout deposit, and of the spatio-temporal variations of the environmental parameters described above.

### 4.2 Physicochemical properties of the respirable volcanic ash

The mineralogical assemblage of the isolated respirable ash sample from the 16-17 August 2006 fallout deposit is characteristic of an andesitic magma (andesitic glass, plagioclase, pyroxene, and Fe-Ti oxides), and is consistent with petrological data acquired on lapilli and bombs from the same eruption.^45^ The other identified phases (Fig. 4, Table 3) are also consistent with the eruptive dynamics of the 2006 Tungurahua eruption:(1) the rhyolitic glass and the sanidine crystals originate from the eruption of a more differentiated magma pocket, as previously demonstrated^45^, and which only represents 0.4 vol.% of the fallout deposit^47^, (2) alunite probably originate from the condensation of volcanic gas on ash particles during their transport in the eruption plume and atmosphere, a well know phenomenon during explosive volcanic eruptions.^65,66^ The origin of the carbonaceous matter is unknown, but can be due to burning vegetation or entrainment of organic material during the eruption. Observation of carbonaceous matter in this respirable volcanic ash further substantiates that people are rarely exposed to volcanic PM alone.^67^

The morphology and surface texture of the respirable ash (Fig. 5) is highly variable and phase dependant. Crystals have the highest surface roughness, and crystal fragments appear not to be broken along grain boundaries (e.g., Fig. 5m). Freshly fractured surfaces are observed on both crystals and glass, and all particle types are covered with sub-1 µm adhering particles, which are predominantly chips of the magmatic particles and, more rarely, plume-derived sulfates (Fig. 5k-n). Overall, the respirable ash analysed here have a low toxic potential, given the absence of mineral phases and particle morphologies of known-pathogenicity, such as crystalline silica and fiber-like particles.^11^

### 4.3 Bioreactivity and implications for the respiratory hazard

Contrary to quartz, exposure to the Tungurahua respirable ash is weakly cytotoxic towards A549 alveolar epithelial type II cells, as demonstrated by the LDH assay (Fig. 7a) and the TEM and SEM imaging (Fig. 6). The SEM images (Fig. 6e-h) demonstrate a change in cell morphology and membrane texture after exposure to respirable ash, which suggests that the cells start undergoing some suffering, and a potential phenotype change. TEM images demonstrate that the respirable ash are internalized by the cells and processed in the endosomal pathway. The fate of these particles after 24 hours is yet to be determined, and the biodurability of ash in intracellular compartments is poorly constrained.^68^ If particles are not efficiently digested, they will remain in the lungs or be translocated, such as to the lymphatic system, as evidenced *in-vivo* via instillation of volcanic ash from the Montserrat Soufrière Hills eruption^16^, where persistence or continued dissolution may result in sustained inflammation.

Despite being a widely used model in toxicology, few studies have explored the response of A549 alveolar epithelial type II cells to respirable volcanic ash. Three studies focused on cytotoxicity using samples from the 18 May 1980 Mount St Helens eruption, USA^69^ and the 1995-2013 Soufrière Hills eruption, Montserrat^70,71^, and only one study also assessed the pro-inflammatory response of these cells (IL-6, IL-8 and IL-1β release) using samples from 6 different volcanic eruptions with variable content of crystalline silica^19^. All of these studies reported little cytotoxicity of volcanic ash, irrespective of the sample, in agreement with our results. Using doses of respirable ash between 5 and 50 µg/cm^2^, and a 24 h timepoint, Damby, et al. ^19^ show that IL-1β is not produced by A549 cells, even following quartz treatment, as observed here even at the higher dose (105 µg/cm^2^). The release of IL-6 and IL-8 observed here after 24 h (Fig. 7c) is not reported by Damby, et al. ^19^, IL-8 not being even produced by quartz treatment in their experiments. These contrasting results could be explained by the difference in particle doses (maximum of 50 µg/cm^2^ in Damby, et al. ^19^ vs. 105 µg/cm^2^ here). The relevance of this difference in maximum dose is unknown given a lack of available *in-vivo* dosimetry data during volcanic eruptions^49^, particularly in the context of the Tungurahua area, given the long exposure duration, spatial variations in the characteristics of volcanic ash deposits and environmental conditions, and disparities in human activity. Regardless, our work demonstrates that respirable volcanic ash from the 16-17 August paroxysmal phase of Tungurahua can initiate a low-level pro-inflammatory response in A549 cells, even when not containing crystalline silica, particularly in the case of chronic exposure. This work also shows that the alveolar epithelium has a weak inflammatory role, but could rather have a signaling role in order to initiate the recruitment of macrophages and the on-set of a global inflammation status.

In the context of the long exposure duration of the populations in the Tungurahua region, these different findings provide justification for dedicated exposure and epidemiological studies. After the first year of activity, increased respiratory infections were reported in Penipe, a town just south of Puela (Fig. 1), which was exposed to the early ashfalls of the Tungurahua eruption but was not evacuated.^43^ Little to no health data on the populations after 2001 have been published, expect for qualitative reports of increased respiratory problems from health care professional described in Sword-Daniels, et al. ^44^. Such reports suggest that a long-term, chronic respiratory hazard could exist, that could be further investigated.

## 5 Conclusion

Health risk assessments at active volcanoes are exceptionally rare, even at highly hazardous volcanoes like Tungurahua, where frequent ash-forming explosive events occurred for almost two-decades, subjecting local populations to a chronic exposure to volcanic PM, and respirable volcanic ash in particular. This study highlights that, in such a context of prolonged volcanic activity, to comprehensively assess the resilience of the Tungurahua social-ecological system, the health risk needs to be included. Indeed, this work evidences that large quantities of respirable volcanic ash was produced by the sole 16-17 August 2006 explosive phase, which affected the whole agricultural area surrounding the volcano, where more than 60% of adults are outdoor workers. We show that PDC and co-PDC formations are key eruptive phenomena contributing significantly to the high content in respirable ash of this explosive phase. Overall, the respirable ash sample has a low toxic potential in view of the mineralogical content and the particle morphologies. The *in-vitro* assays on A549 cells, a widely used model for human alveolar epithelial type II cells, demonstrate that respirable volcanic ash is efficiently internalized by the cells and processed in the endosomal pathway, and no membrane destabilization is observed, which should be further investigated to understand the fate of these particles after 24 hours. Additionally, the acute inflammatory response of these cells evidences the potential for the Tungurahua respirable ash to initiate a low-level inflammatory signal, which could lead to chronic inflammation by recruitment of immune cells. Further research needs to be carried out to understand the biological response to a chronic exposure to this volcanic ash.

This work provides the first mapping of inhalable and respirable volcanic ash around an active volcano, and details a systematic and large spatial variability, a critical parameter to account for during a respiratory health hazard assessment and interpretations of exposures. These novel results provide grounds for future exposure measurements and air quality monitoring campaigns, particularly targeting outdoor workers. We also emphasize the importance of using the mass per unit area of inhalable volcanic ash deposited for quantifying the respiratory health hazard. During the Tungurahua eruption, the amount of respirable ash as well as their physicochemical properties are dependent on the eruption dynamics (e.g., eruptive phenomena such as formation of PDCs, mechanisms of ash transport and deposition). Quantifying the eruption dynamics is thus of critical importance for health hazard assessment.

## Acknowledgments

J.E., L.G., V.S. and L.B. acknowledge funding from the University Clermont-Auvergne I-Site CAP20-25 program that supported this work. J.E. thanks Jeanne Tran Van Nhieu and Pascale Gueirard for invaluable insights on the TEM images. D.D. acknowledges support from the WOW! Visiting Scholar Fellowship from the University Clermont-Auvergne. Any use of trade, firm, or product names is for descriptive purposes only and does not imply endorsement by the U.S. Government. This is Laboratory of Excellence ClerVolc contribution n°XXX.

