## Supplemental Material for "Spatial distribution and physicochemical properties of respirable volcanic ash from the 16-17 August 2006 Tungurahua eruption (Ecuador), and alveolar epithelium response *in-vitro*"

Julia Eychenne[1,2*], Lucia Gurioli[1], David Damby[3], Corinne Belville[2], Federica Schiavi[1], Geoffroy Marceau[2,4], Claire Szczepaniak[5], Christelle Blavignac[5], Mickael Laumonier[1], Jean-Luc Le Pennec[6,7], Jean-Marie Nedelec[8], Loïc Blanchon[2], Vincent Sapin[2,4]

*[1] Université Clermont Auvergne, CNRS, IRD, OPGC, Laboratoire Magmas et Volcans, F-63000 Clermont-Ferrand, France*

*[2] Université Clermont Auvergne, CNRS, INSERM, Institut de Génétique Reproduction et Développement, F-63000 Clermont-Ferrand, France*

*[3] U.S. Geological Survey, California Volcano Observatory, Moffett Field, CA, USA*

*[4] Biochemistry and Molecular Genetic Department, University Hospital, F-63000 Clermont-Ferrand, France*

*[5] Université Clermont Auvergne, UCA PARTNER, Centre Imagerie Cellulaire Santé, F-63000 Clermont-Ferrand, France*

*[6] Geo-Ocean, CNRS, Ifremer, UMR6538, F-29280 Plouzané, France*

*[7] IRD Office for Indonesia & Timor Leste, Jalan Kemang Raya n°4, Jakarta 12730, Indonesia*

*[8] Université Clermont Auvergne, SIGMA Clermont, F-63000 Clermont-Ferrand, France*

### Content of this file:

**Supplemental Figures S1 to S2 with captions**

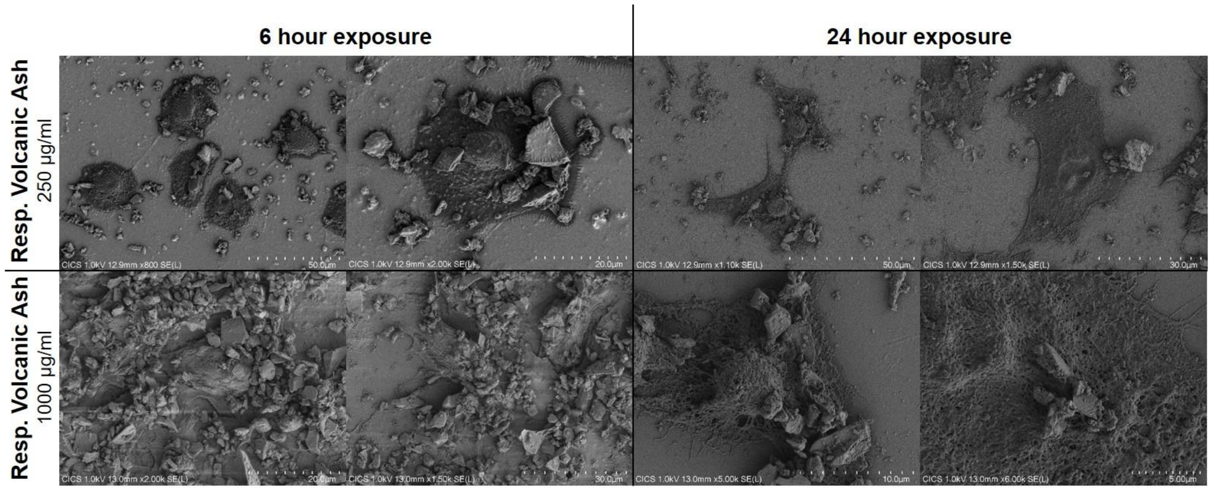


**Fig. S1:** Interactions between A549 cells and respirable Tungurahua volcanic ash at doses of 250 µg/ml and 1000 µg/ml after 6 and 24h of exposure, imaged by FEG-SEM.

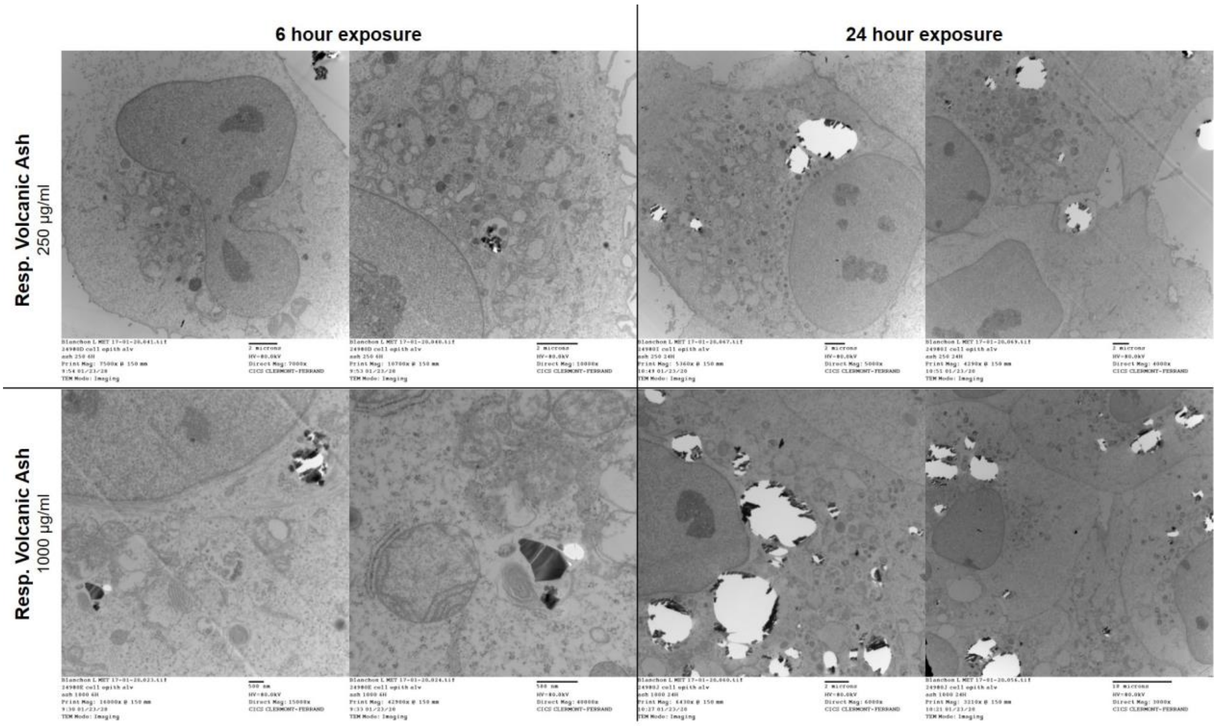


**Fig. S2:** Interactions between A549 cells and respirable Tungurahua volcanic ash at doses of 250 µg/ml and 1000 µg/ml after 6 and 24h of exposure, imaged by TEM.
